# Modeling large fluctuations of thousands of clones during hematopoiesis: the role of stem cell self-renewal and bursty progenitor dynamics in rhesus macaque

**DOI:** 10.1101/343160

**Authors:** Song Xu, Sanggu Kim, Irvin S. Y. Chen, Tom Chou

## Abstract

In a recent clone-tracking experiment, millions of uniquely tagged hematopoietic stem cells (HSCs) were autologously transplanted into rhesus macaques and peripheral blood containing thousands of tags were sampled and sequenced over 14 years to quantify the abundance of hundreds to thousands of tags or “clones.” Two major puzzles of the data have been observed: consistent differences and massive temporal fluctuations of clone populations. The large sample-to-sample variability can lead clones to occasionally go “extinct” but “resurrect” themselves in subsequent samples. Although heterogeneity in HSC differentiation rates, potentially due to tagging, and random sampling of the animals’ blood and cellular demographic stochasticity might be invoked to explain these features, we show that random sampling cannot explain the magnitude of the temporal fluctuations. Moreover, we show through simpler *neutral* mechanistic and statistical models of hematopoiesis of tagged cells that a broad distribution in clone sizes can arise from stochastic HSC self-renewal instead of tag-induced heterogeneity. The very large clone population fluctuations that often lead to extinctions and resurrections can be naturally explained by a generation-limited proliferation constraint on the progenitor cells. This constraint leads to bursty cell population dynamics underlying the large temporal fluctuations. We analyzed experimental clone abundance data using a new statistic that counts clonal disappearances and provide least-squares estimates of two key model parameters in our model, the total HSC differentiation rate and the maximum number of progenitor-cell divisions.

**Author summary:** Hematopoiesis of virally tagged cells in rhesus macaques is analyzed in the context of a mechanistic and statistical model. We find that the clone size distribution and the temporal variability in the abundance of each clone (viral tag) in peripheral blood are consistent with (i) stochastic HSC self-renewal during bone marrow repair, (ii) clonal aging that restricts the number of generations of progenitor cells, and (iii) infrequent and small-size samples. By fitting data, we infer two key parameters that control the level of fluctuations of clone sizes in our model: the total HSC differentiation rate and the maximum proliferation capacity of progenitor cells. Our analysis provides insight into the mechanisms of hematopoiesis and a framework to guide future multiclone barcoding/lineage tracking measurements.

## Introduction

Hematopoiesis is a process by which hematopoietic stem cells (HSCs) produce all the mature blood in an animal through a series of proliferating and differentiating divisions [1]. Maintenance of balanced hematopoietic output is critical for an organism’s survival and determines its response to disease and clinical procedures such as bone marrow transplantation [2–5]. How the relatively small HSC population generates more than 10^11^ cells of multiple types daily over an organism’s lifetime has yet to be fully understood. HSCs are defined primarily by their function but are often quiescent [6]. *In vivo*, it is hard to track the dynamics of individual HSCs, while HSCs *in vitro* do not typically proliferate or differentiate as efficiently. Therefore, the dynamics of HSCs can be inferred only from analyses of populations of progenitors and differentiated blood cells [7] and it is useful to investigate HSC dynamics through mathematical modeling and simulations [8–10].

While most studies model population-level HSC behavior [5, 11, 12], certain aspects of HSCs, such as individual-level heterogeneity in repopulation and differentiation dynamics, have to be studied on a single-cell or clonal level [13]. Single HSC transplant mouse data [14] and clonal tracking of HSCs [15,16] in mice have shed some light on repopulation dynamics under homeostasis and after bone marrow transplantation [5,17,18]. However, murine studies usually involve only one or a few clones. How each individual HSC contributes to the blood production process over long times in much larger human and non-human primates is less clear and more difficult to study. Also, unlike in mice, there is no way to isolate and mark HSC populations in human [19].

Recently, results of a long-term clonal tracking of hematopoiesis in normal-state rhesus macaques has been made available [13, 20]. The experiment extracted and uniquely “labelled” hematopoietic stem and progenitor cells (HSPCs) from four rhesus macaques with viral tags that also carry an enhanced green fluorescent protein gene. After autologous transplantation, if any of the tagged HSPCs divide and differentiate, its progeny will inherit their unique tags and ultimately appear in the peripheral blood. Blood samples were drawn every few months over 4 − 14 years (depending on the animal) and the sampled cells were counted and sequenced. Of the ~ 10^6^ − 10^7^ unique HSC tags transplanted, ~ 10^2^ − 10^3^ clones were detected in the sampled peripheral blood. In the original paper describing the clonal tracking experiment, Kim *et al.* [13] observed “A small fraction (4−10%) of tagged clones predominately contribute to a large fraction (25 − 71%) of total blood repopulation.” They described the fluctuations of tags that appeared in each sample as “waves of clones”, but did not address why some clones can disappear at certain times and reappear in a latter sample.

In this study, we seek to better understand the observed clone size distributions and the large temporal variability in clonal populations. To address these observations, we ask: Is heterogeneity in HSCs necessary for peripheral blood clone size heterogeneity, or can a neutral model explain clone size differences? Are clones that disappear and reappear from sample to sample simply missed by random blood sampling, or do other mechanisms of temporal variability need to be invoked?

Unlike other previous models that describe the evolution of lineages of different cell types and their regulation [8–10,21], we will consider simpler neutral models that describe the dynamics of specifically granulocyte populations carrying different tags. Of central interest is the competition among the thousands of clones under a neutral environment that gives rise to fluctuations, extinctions, and resurrections in individual clone populations. Even when considering only one cell type, realistic mathematical models may need to include complex multilevel biochemical feedback mechanisms of regulation [8,22–27]. Many mechanisms may contribute to temporal fluctuations, including extrinsic noise and heterogeneity of HSCs, progenitors, or mature granulocytes. Large time gaps between samplings (5 − 11 months) and small sample sizes also add to the uncertainty of the underlying dynamics. Trying to infer all possible mechanisms and associated parameters from the experimental data would essentially be an overfitting problem. In order to feasibly compare with experimental data, our modeling philosophy will be to recapitulate these complexities into simple, effective models and infer parameters that subsume some of these regulatory effects. This approach and level of modeling are similar to those taken by *e.g.,* Yang, Sun, and Komarova [28, 29].

After careful consideration of a number of key physiological mechanisms, we hypothesize that stochastic HSC self-renewal, generation-limited progenitor cell proliferation, and sampling frequency statistics provide the simplest reasonable explanation for the observed clonal size variability and large temporal fluctuations. HSCs that are generated from self-renewal of the founder population share the same tag as their founder HSC. Thus, during intense self-renewal after myeloablative treatment and HSPC transplantation, each originally transplanted HSCs begets a clonal HSC subpopulation. Subsequently, heterogeneous clone sizes are stochastically generated even though each tag was initially represented by only a single cell. These expanded HSC clones then go on to repopulate the clones in the progenitor and mature blood population, which are also distinguishable by their corresponding tags.

Relative to HSCs, progenitor cells have limited proliferative potential that can explain the apparent extinctions of clones. This limited proliferation potential can be thought of as an “aging” process. Different types of aging, including organism aging [23, 30, 31], replicative senescence of stem cells [32], and generation-dependent birth and death rates, have been summarized by Edelstein *et al.* [33]. Here, the clonal “aging” mechanism we invoke imposes a limit to the number of generations that can descend from each newly created (from HSC differentiation) “zeroth generation” progenitor cell. Possible sources of such a limit include differentiation-induced loss of division potential [34] and telomere shortening (as in the Hayflick limit) [35–37]. Mathematically, genealogical aging can be described by tracking cell populations within each generation. After a certain number of generations, progenitor cells of the final generation stop proliferating and can only differentiate into circulating mature cells or die.

In the following sections, we first present the mathematical equations and corresponding solutions (whenever possible) of a model that incorporates the above processes. We then develop a new statistical measure that tracks the numbers of absences of clones across the samples. Measured clone abundances of animal RQ5427 are statistically analyzed within our mechanistic model to infer estimates for key model parameters. The data and corresponding statistical analyses for animals 2RC003 and RQ3570 are also provided in the Results section.

## Materials and Methods

Below, we describe available clonal abundance data, mechanistic models, and a statistical model we will use for parameter inference.

### Clone abundance data

In the experiments of Kim *et al.* [13], cells in samples of peripheral blood were counted to extract 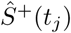, the total number of EGFP+ tagged cells in sample 1 ≤ *j* ≤ *J* taken at time *t_j_*. After PCR amplification and sequencing, 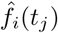, the relative abundance of the *i*^th^ tag among all sampled, tagged cells is also quantified. The “ˆ” notation will henceforth indicate experimentally measured quantities.

Within mature peripheral blood, lymphocytes such as T cells and B cells proliferate or transform in response to unpredictable but clone-specific immune signals [38]. They also vary greatly in their lifespans, ranging from days in the case of regular T and B cells to years in the case of memory B cells. On the other hand, mature granulocytes do not proliferate in peripheral blood and have relatively shorter life spans [7]. Granulocyte dynamics can thus be analyzed with fewer confounding factors [11]. Thus, in this paper, we restrict our analysis to granulocyte repopulation and extract all variables, including 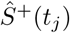 and 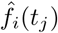 described above, that are associated exclusively with granulocyte populations.

In Fig. 1(a), we plot the total numbers of sampled granulocytes from one of the macaques, RQ5427. The subpopulation of EGFP+ granulocytes and the subset of EGFP+ granulocytes that were extracted for PCR amplification and analysis are also plotted. Data for two other animals, 2RC003 and RQ3570, are qualitatively similar, while those for the fourth animal, 95E132, did not separate granulocytes from peripheral blood mononuclear cells. As shown in Fig. 1(b), not only are the clone abundances 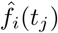 heterogeneous, but individual clone abundances vary across samples taken at different times. The variation is so large that many clones can go extinct and reappear from one sample to another, as shown in Fig. 1(c). Since large numbers of progenitor and mature cells are involved in blood production, the observed clone size fluctuations cannot arise from intrinsic demographic stochasticity of progenitor- and mature-cell birth and death. Moreover, we will show later in the Results section that random sampling alone cannot explain the observed clonal variances and mechanisms that involve other sources of variation are required.

**Fig 1.**
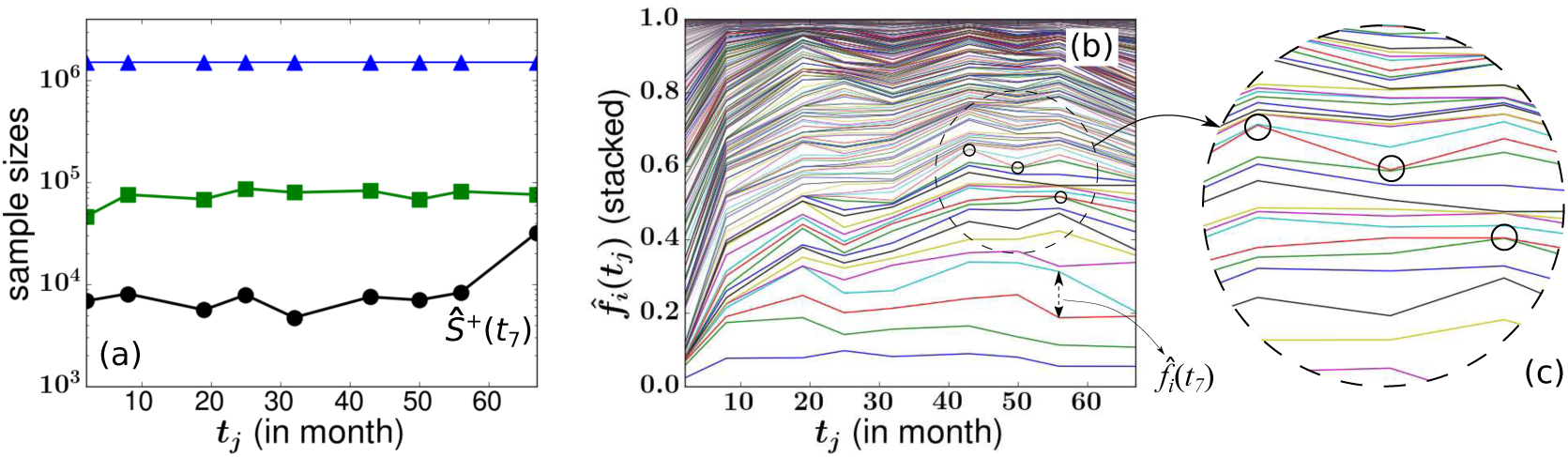
Blood sample data from animal RQ5427 [13]. (a) The total numbers of sampled granulocytes (blue triangles), EGFP+ granulocytes (green squares), and the subset of EGFP+ granulocytes that were properly tagged and quantifiable were extracted for PCR amplification and analysis (black circles). This last population defined by 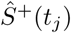 is used to normalize clone cell counts. We excluded the first sample at month 2 in our subsequent analysis so, for example, the sample at month 56 is labeled the 7th sample. There were 536 clones detected at least once across the eight samples taken over 67 months comprising an average fraction 0.052 of all granulocytes. The abundances of granulocyte clones are shown in (b). The relative abundance 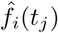 of granulocytes from the *i*^th^ clone measured at month *t_j_* is indicated by the vertical distances between two adjacent curves. The relative abundances of individual clones feature large fluctuations over time. “Extinctions” followed by subsequent “resurrections,” were constantly seen in certain clones as indicated by the black circles in (b) and in the inset (c).

### Nomenclature and lumped mechanistic model

Fig. 2 depicts our neutral model of hematopoiesis which is composed of five successive stages, or compartments, describing the initial single-cell tagged HSC clonal populations immediately after transplantation (Compartment **0**), the heterogeneous HSC clonal populations after a short period of intense self-renewal (Compartment **1**), the transit-amplifying progenitor cell compartment (Compartment **2**), the peripheral blood pool (Compartment **3**), and the sampled peripheral blood (Compartment **4**), respectively. Each distinct color or shape in Fig. 2 represents a distinct clone of cells with the same tag.

**Fig 2.**
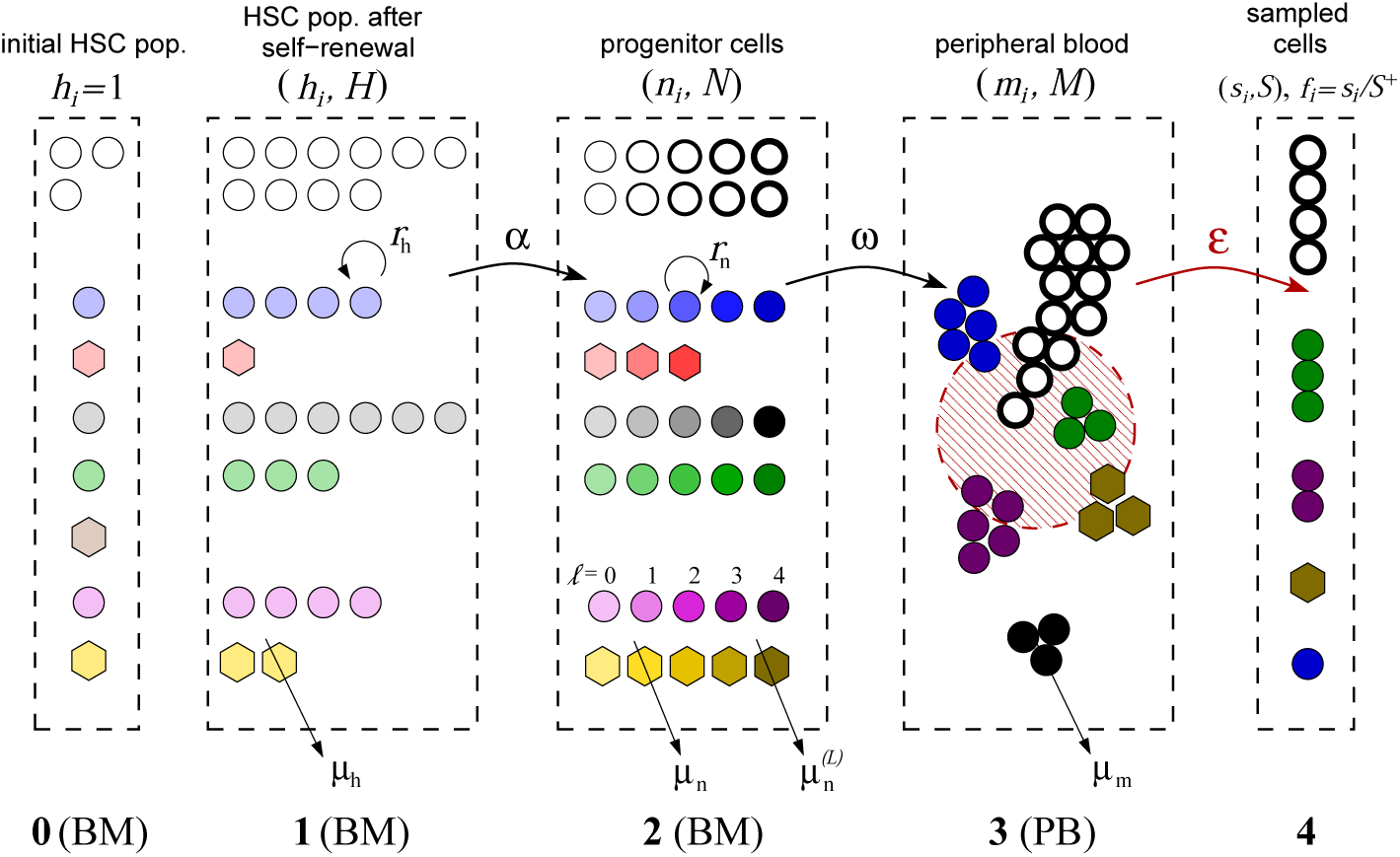
Schematic of a neutral multi-stage or multi-compartment hematopoiesis model. BM and PB refer to bone marrow and peripheral blood, respectively. Cells of the same clone have the same color. White circles represent untagged cells which were not counted in the analysis. Stages **0**, **1**, and **2** describe cell dynamics that occur mainly in the bone marrow. Stage **1** describes HSC clones (*C*_h_ = 6 in this example) after self-renewal that starts shortly after transplantation with rate *r*_h_. After self-renewal, the relatively stable HSC population (*H*^+^ = 20 in this example) shifts its emphasis to differentiation (with per-cell differentiation rate *α*). Larger clones in Stage **1** (*e.g.*, the circular blue clone, *h*_blue_ = 4) will have a larger total differentiation rate *αh*_blue_ while smaller clones (*e.g.*, the red hexagonal clone, *h*_red_ = 1) will have smaller *αh*_red_. The processes of progenitor-cell proliferation (with rate *r*_n_) and maturation (with rate *ω*) in Compartments **2** and **3** are considered deterministic because of the large numbers of cells involved. The darker-colored symbols correspond to cells of later generations. For illustration, the maximum number of progenitor-cell generations allowed is taken to be *L* = 4. Compartment **4** represents a small sampled fraction (*ε*(*t_j_*) ≈ 2.8 × 10^−5^ − 2 × 10^−4^) of Compartment **3**, the entire peripheral blood of the animal. In the example pictured above, *C*_s_ = 4. Such small samples can lead to considerable sampling noise but is not the key driver of sample-to-sample variability.

In each compartment, relevant parameters include (using Compartment **1** as example): the total cell count *H*(*t*), the untagged cell count *H*^−^(*t*), the tagged cell count *H*^+^(*t*), the total number of tagged clones *C*_h_(*t*), and the number *h_i_*(*t*) of HSCs carrying the *i*^th^ tag. These quantities are related through 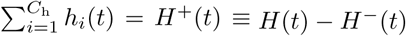.

In the progenitor pool, the total number of cells and the number with tag *i* are denoted *N*(*t*) and *n_i_*(*t*), respectively. Further resolving these progenitor populations into those of the *ℓ*^th^ generation, we define *N*^(*ℓ*)^(*t*) and 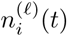. In the mature granulocyte pool, the total granulocyte population and that with tag *i* are labelled *M*(*t*) and *m_i_*(*t*). In the sampled blood compartment, we use *S*(*t_j_*), *S^+^*(*t_j_*), *s_i_*(*t_j_*), and *C_s_*(*t_j_*) to denote, at time *t_j_*, the total number of sampled cells, the number of tagged sampled cells, the total number of tagged cells of clone *i*, and the total number of clones in the sample, respectively. In Compartment 4, we further define *f_i_*(*t_j_*) = *s_i_*(*t_j_*)*/S^+^*(*t_j_*) to denote the relative abundance of the *i*^th^ clone among all tagged clones.

By lumping together all clones (tagged and untagged) in each compartment, we can readily model the dynamics of total populations in each pool. After myeloablative treatment, the number of BM cells, including HSCs, is severely reduced. Repopulation of autogolously transplanted HSCs occurs quickly via self-renewal until their total number *H*(*t*) reaches a steady-state. The repopulation of the *entire* HSC population and the subsequent entire progenitor and mature cell populations may be described via simple deterministic mass-action growth laws

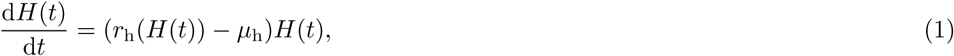

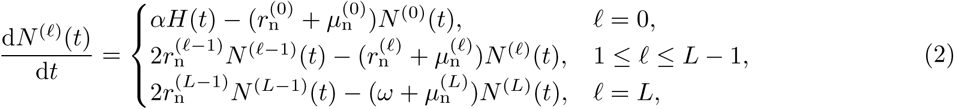

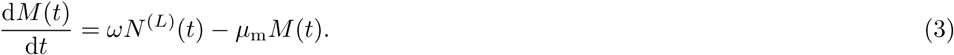

HSC self-renewal is a regulated process involving signaling and feedback [22–24,39,40] and *r*_h_ may be a complicated function of many factors; however, we will subsume this complexity into a simple population-dependent logistic law *r*_h_(*H*(*t*)) and assume a constant death rate *µ*_h_. Alternatively, other studies have employed Hill-type growth functions [12, 28].

We assume the per cell HSC differentiation rate *α* is independent of the tag and that differentiation is predominantly an asymmetric process by which an HSC divides into one identical HSC and one progenitor cell that commits to differentiation into granulocytes. An initial generation-zero progenitor cell further proliferates with rate 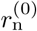, contributing to the overall progenitor-cell population. Subsequent generation-*ℓ* progenitors, with population *N*^(*ℓ*)^, proliferate with rate 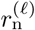 until a maximum number of generations *L* is reached. By keeping track of the generation index *ℓ* of any progenitor cell, we limit the proliferation potential associated with an HSC differentiation event by requiring that any progenitor cell of the final *L*^th^ generation to terminally differentiate into peripheral blood cells with rate *ω* or to die with rate 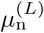. For simplicity, we neglect any other source of regulation and assume *α*, 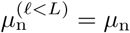, 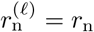 and *ω* are all unregulated constants.

Our model analysis and data fitting will be performed using clone abundances sampled a few months after transplantation under the assumption that granulopoiesis in the animals has reached steady-state [4] after initial intensive HSC self-renewal. Steady-state solutions of Eqs. (1), (2) and (3) are defined by *H_ss_*, 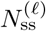, and *M_ss_*. The first constraint our model provides relates these steady-state populations through

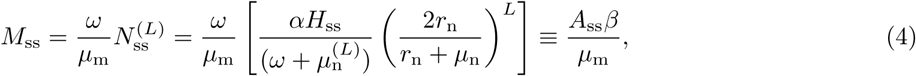

where we have defined

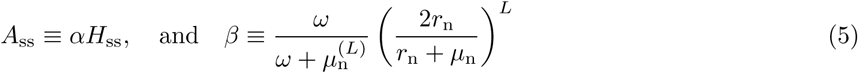

as the total rate of HSC differentiation and the average number of granulocytes generated per HSC differentiation, respectively. These constraints also hold for the EGFP+ subset (about 5%-10%) of cells, *e.g.*, 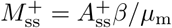 and 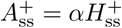. Since 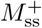 is inferred from the experiment, Eq. (4) places a constraint between 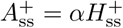 and *β*. This steady-state constraint will eventually be combined with statistics of the fluctuating clone abundances data to infer estimates for the underlying model parameters.

### Clone-resolved mechanistic model

Although the lumped model above provides important constraints among the steady-state populations within each compartment, the clone-tracking experiment keeps track of the populations of sampled granulocytes that arise from “founder” HSCs that carry the same tag. Thus, we need to resolve the lumped model into the clonal subpopulations described by *h_i_*, 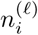, and *m_i_*.

Even though the total HSC populations *H*(*t*) and *H*^±^(*t*) are large, the total number of clones *C*_h_ ≫ 1 in compartment 1 is also large, and the number of cells with any tag (the size of any clone) can be small. The population of cells with any specific tag *i* is thus subject to large demographic fluctuations. Thus, we model the stochastic population of HSCs of any tag using a master equation for *P*(*h*, *t*), the probability that at time *t* the number of HSCs of any clone is *h:*

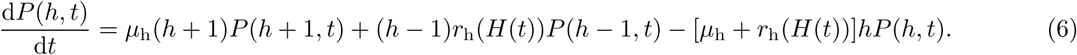

Recall that immediately after transplantation, each HSC carries a distinct tag before self-renewal (*h_i_*(0) = 1) leading to the initial condition 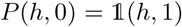, where the indicator function 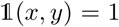 if and only if *x* = *y*. Because *h* = 0 is an absorbing boundary, clones start to disappear at long times resulting in a decrease in the total number *C*_h_(*t*) of HSC clones. Before this “coarsening” process significantly depletes the entire population, each clone constitutes a small subpopulation among all EGFP+ cells, *h*(*t*) ≪ *H*(*t*), and the stochastic dynamics of the population *h* of any clone can be approximated by the solution to Eq. (6) with *r*_h_(*H*(*t*)) replaced by *r*_h_(*t*). Hence, evolution of each HSC clone follows a generalized birth-death process with time-dependent birth rate and constant death rate. We show in Appendix A in S1 Text that for *H* ≫ 1 the solution to Eq. (6) can be written in the form [41]

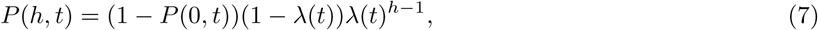

where 0 ≤ *λ*(*t*) < 1 depends on *r*_h_(*t*) and *µ*_h_. Here, *λ*(*t*) determines “broadness” (level of clone size heterogeneity) of the clone size distribution. For the relevant initial condition of unique tags at *t* = 0, *λ*(0) = 0 and *λ*(*t* → ∞) → 1. When *λ*(*t*) is small, the distribution is weighted towards small *h*. For *λ*(*t*) = 0, 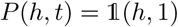 which was the limit used in Goyal *et al.* [4] to assume no HSC self-renewal after transplantation. In the limit *λ*(*t*) → 1, the distribution becomes flat and a clone is equally likely to be of any size 1 ≤ *h* ≤ *H*.

To further resolve the progenitor population into cells with distinct tags, we define *n*^(*ℓ*)^ (*t*) as the number of generation-*ℓ* progenitor cells carrying any one of the viral tags. The total number of progenitor cells with a specific tag is 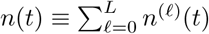. Since the sizes *h_i_* of individual clones may be small, differentiation of HSCs within each clone may be rare. However, since the size of each tagged progenitor clone quickly becomes large (*n*(*t*) ≫ 1), we model the dynamics of *n*^(*ℓ*)^ (*t*) using deterministic mass-action growth laws:

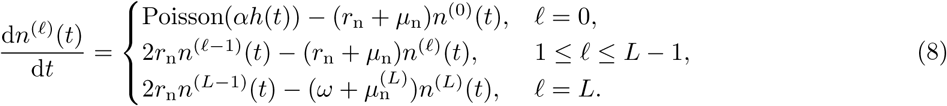

Our model is neutral (all clones have the same birth, death, and maturation rates), so these equations are identical to Eqs. (2). However, since creation of the zeroth-generation subpopulation *n*^(0)^ (*t*) derives only from differentiation of HSCs of the corresponding clone, which has a relatively small population *h*(*t*), we invoke a Poisson process with rate *αh*(*t*) to describe stochastic “injection” events associated with asymmetric differentiation of HSCs of said clone. Each discrete differentiation event leads to a temporal burst in *n*^(*ℓ*)^(*t*).

Finally, the dynamics of the population *m*(*t*) of any granulocyte clone in the peripheral blood are described by an equation analogous to Eq. (3):

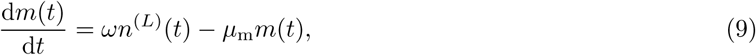

where we have assumed that only the generation-*L* progenitor cells undergo terminal differentiation with rate *ω*. An alternative model allows progenitor cells of earlier generations (*ℓ* < *L*) to also differentiate and circulate but does not give rise to qualitatively different results (See Appendix B in S1 Text).

To study the dynamics of the burst in 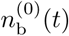 immediately following a *single*, *isolated* asymmetric HSC differentiation event at *t* = 0, we set the initial condition 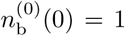, 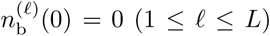, remove the Poisson(*αh*(*t*)) term in Eq. (8) and find,

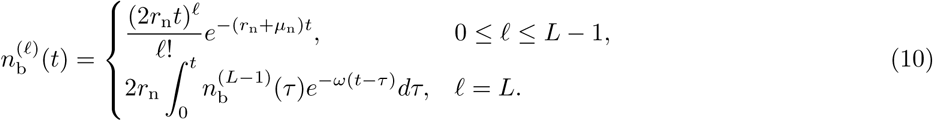

Bounded analytic solutions to 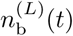 involving the lower incomplete gamma function can be found. Upon using the solution 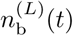 (t) in Eq. (9) the mature blood population within a clone associated with a single HSC clone differentiation even is described by

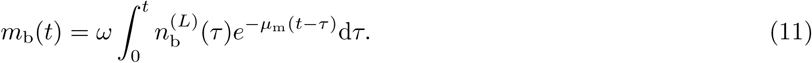

The populations associated with a single HSC differentiation event, 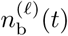 and 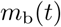, are plotted below in Fig. 3 of the Results section. Then *m_i_*(*t*), the total number of mature cells with the *i*^th^ tag at time *t*, is obtained by summing up all *m*_b_(*t* − *τ_k_*) bursts initiated by HSC differentiations at separate times *τ_k_* ≤ *t* with the *i*^th^ tag.

**Fig 3.**
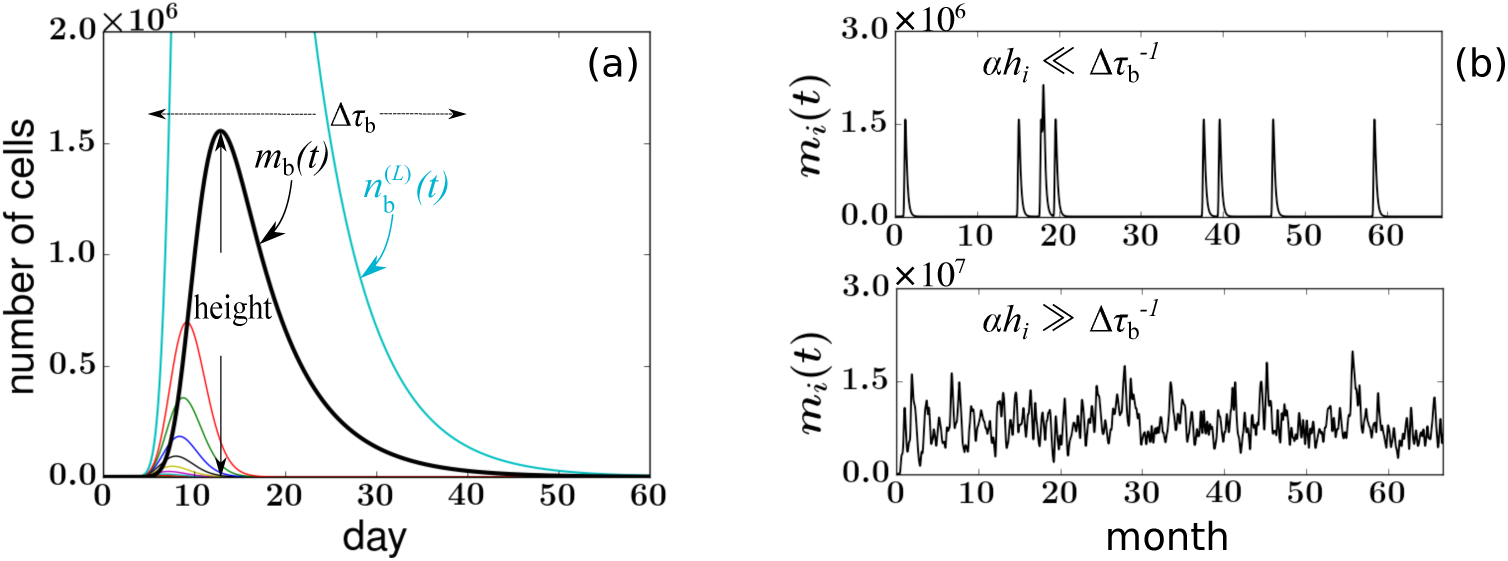
(a) A burst of cells is triggered by a single HSC differentiation event at time *t* = 0. A plot of representative solutions to Eqs. (10) and (11) for *r*_n_ = 2.5, *L* = 24, 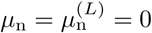, 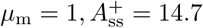, and *ω* = 0.16. Curves of different colors represent 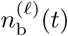, the progenitor cell population within each generation *ℓ* = 0, 1, 2, …, *L*, and *m*_b_(*t*), the number of mature granulocytes associated with the differentiation burst. All populations rise and fall. (b) Realizations of PB numbers of a single clone arising from multiple successive differentiation events. The fluctuating populations are generated by adding together *m*_b_(*t*) associated with each differentiation event. Time series resulting from small (*h_i_*/*H*^+^ = 0.0003) and large (*h_i_*/*H*^+^ = 0.03) HSC clones are shown. Small clones are characterized by separated bursts of cells, after which the clone vanishes for a relatively long period of time. The number of mature peripheral blood cells of large clones reaches a relatively constant level and almost never vanishes.

Besides the burst dynamics described above, the data shown in Fig. 1(a) are subject to the effects of small sampling size, uncertainty, and bias induced by experimental processing such as PCR amplification, and data filtering. In this experimental system, PCR generates a smaller uncertainty than blood sampling so we focus on the statistics of random sampling. Each blood sample drawn from monkey RQ5427 contains about 10µg of genomic DNA [13]. After PCR amplification, deep sequencing, and data filtering, the total number 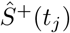 of quantifiable tags corresponds to ~ 5 × 10^3^ − 3 × 10^4^ tagged cells. The sample ratio is defined by 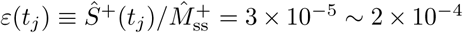 where 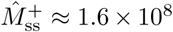 is the estimated total number of tagged cells in the peripheral blood. The number of sampled cells with the *i*^th^ tag from the *j*^th^ sample then approximately follows a Binomial distribution 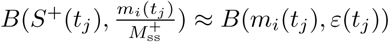 in our model. To quantitatively explore the feature of apparent extinctions of clones from a sample, we calculate the probability that no peripheral blood cell from clone *i* is found in a sample of size 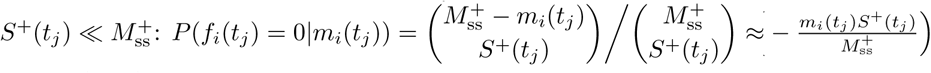. Thus, if 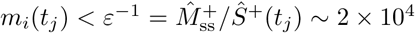 the *i*^th^ clone is likely to be missed in the sample. The value *ε*^−1^ is also used to threshold the population *m*_b_(*t*) to define the measurable duration Δ*τ*_b_ of a burst (as indicated in Fig. 3(a)).

### Parameter values

Parameters determined by the experimental procedure or estimated directly from the experiments include the weight of the animal, the sampling times *t_j_*, the EGFP+ ratio, and the total number of tagged cells detected in each sample 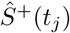. Since 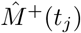 does not fluctuate much, we use its average for 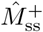 and the relevant experimental parameters for each animal become 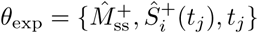. These will also be used as inputs to our models.

Our multi-stage model also contains many other intrinsic parameters, including 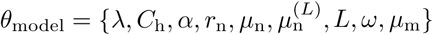. We first found parameter values that have been reliably independently measured. Some parameters were measured in human clinical studies rather than in rhesus macaques, but can nonetheless serve as reasonable approximations for non-human primates due to multiple physiological similarities [42]. These estimates can certainly be improved once direct measurements on rhesus macaques become available. Model parameters, their estimates, and the associated references are given in Table 1 below.

**Table 1.** Summary of parameters, including their biological interpretation, ranges of values, and references. All rate parameters are quoted in units of per day. Other parameters are chosen to be within their corresponding reported ranges from the referenced literature. How variations in parameter values of affect our analysis will be described in the subsequent sections.

| Parameter | Interpretation | Values & References |
| --- | --- | --- |
| **HSC pool (Compartments 1)** |  |  |
| $H_{ss}$ | total number of HSCs at steady state | $1.1 \times 10^4 - 1.1 \times 10^6$ [4, 11, 12] |
| $\alpha$ | per-cell HSC differentiation rate | $5.6 \times 10^{-4} - 0.02$ [4, 11, 12] |
| $\mu_h$ | HSC death rate | $10^{-3} - 0.1$ [12, 34] |
| **Transit Amplifying Progenitor pool (Compartment 2)** |  |  |
| $r_n$ | growth rate of progenitor cell | $2 - 3$ [12] |
| $\mu_n$ | death rate of progenitor cell (generation $\ell < L$) | $0$ [12, 34] |
| $\mu_n^{(L)}$ | death rate of progenitor cell (generation $\ell = L$) | $0 - 0.27$ [12, 34] |
| $\omega$ | maturation rate of generation-$L$ cells | $0.15 - 0.17$ [43, 44] |
| $L$ | maximum generation of progenitor cells | $15 - 21$ [12, 34] |
| **Peripheral Blood pool (Compartment 3)** |  |  |
| $M_{ss}$ | total number of peripheral blood granulocytes at steady state | $(2.5 - 5) \times 10^9$ [13, 42] |
| $\mu_m$ | death rate of peripheral blood granulocytes | $0.2 - 2$ [34, 44, 45] |

### Model properties and implementation

Using parameter estimates, we summarize the dynamical properties of our model and describe how the key model ingredients including stability of HSC clone distributions and subsequent “bursty” clone dynamics that follow differentiation can qualitatively generate the observed clone-size variances.

#### Slow homeostatic birth-death of HSCs

The first important feature to note is the slow homeostatic birth-death of HSCs. After the bone marrow is quickly repopulated, *r*_h_(*H*(*t*)) − *µ*_h_ ≈ 0, and stochastic self-renewal slows down. Because *h* = 0 is an absorbing state, the size distribution of the clones may still slowly evolve and coarsen due to stochastic dynamics leading to the slow successive extinction of smaller clones. The typical timescale for overall changes in *h* can be estimated by approximating *r*_h_(*H*_ss_) ≈ *µ*_h_ [46] and considering the mean time *T*(*h*) of extinction of a clone initially at size *h* ≪ *H*_ss_. The standard result given in Gardiner [47] and also derived in Appendix C in S1 Text is 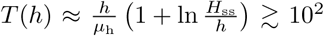 months (for *µ*_h_ = 10^−2^, *H*_ss_ = 10^4^, *h* = 10^1^; see Table 1 for applicable values). Since this timescale is larger than the time of the experiment (67 months for monkey RQ5427), mean HSC clone sizes do not change dramatically during the experiment, consistent with the stable number of clones observed in the samples (for monkey RQ5427, the number of detected clones at month {2, 8, 19, 25, 32, 43, 50, 56, 67} are *C*_s_(*t_j_*) = {184,145,186,193,152,189,155, 286}) shown in Fig. 1(b). Thus, as a first approximation, we will use a static configuration {*h_i_*} drawn from *P*(*h*) to describe how, through differentiation, HSC clones feed the progenitor pool.

#### Fast clonal aging of progenitors

In contrast to slow HSC coarsening, progenitor cells proliferate “transiently.” We plot a single burst of progenitor and mature granulocytes, Eqs. (10) and (11), in Fig. 3(a) using the parameter values listed in Table 1. Associated with each temporal burst of cells, we define the characteristic duration, or “width” Δ*τ*_b_ as the length of time during which the number *m*_b_(*t*) is above the detection threshold within a sample of peripheral blood: 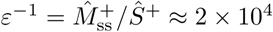.

According to Eq. (11), the burst width and height depend nonlinearly on the parameters *L*, *r*_n_, *µ*_n_, *µ*_m_, and *ω* in their physiological ranges (see Table 1). The characteristic “width” of a burst scales as Δ*τ*_b_ ~ *L*/*r*_n_ + *1*/*ω* + 1/*µ*_m_. This estimate is derived by considering the *L* rounds of progenitor cell division, each of which takes time ~ 1/*r*_n_. Terminal-generation progenitors then require time ~ 1/*ω* to mature, after which mature granulocytes live for time ~ 1/*µ*_m_. In total, the expected life span of ~ *L/r*_n_ + 1/*ω* + 1/*µ*_m_, which approximates the timescale of a HSC-differentiation-induced burst of cells fated to be granulocytes. Using realistic parameter values, the typical detectable burst duration Δ*τ*_b_ ~ 1 − 2 months is much shorter than the typical sampling gaps Δ*t_j_* = 5 − 11 months.

With this “burst” picture in mind, we now show how fluctuations of sampled clone sizes can be explained. Small-*h* 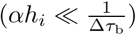 clones never or rarely appear in blood samples. Their appearance also depends on whether sampling is frequent and sensitive enough to catch the burst of cells after rare HSC differentiation events. On the other hand, large-*h* 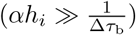 clones differentiate frequently and consistently appear in the peripheral blood. Their populations in blood samples are less sensitive to the frequency of taking samples. Fig. 3(b) shows two multi-burst realizations of *m_i_*(*t*) corresponding to two values of *h_i_*. The 2000-day trajectories were simulated by fixing *h_i_* and stochastically initiating the progenitor proliferation process. Population bursts described by Eq. (11) were added after each differentiation event distributed according to Poisson(*αh_i_*).

Thus, the statistics of clone extinctions and resurrections should be more sensitive to the overall clonal differentiation rate *αh_i_* than to the precise shape of a mature cell burst. This is confirmed by further simulation studies and analysis (Appendix D in S1 Text) and motivates reducing the number of effective parameters.

We can further pare down the number of remaining parameters by finding common dependences in the model and defining an effective maximum generation number. We can rewrite Eq. (5) as 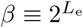, where

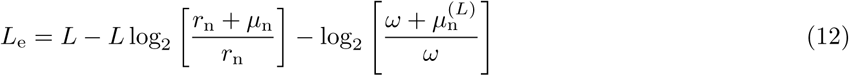

is an *effective* (and noninteger) maximum generation parameter. Later in Appendix D in S1 Text, we show that uncertainties of the model structure, alternative mechanisms, and parameter values can be subsumed into *L*_e_. Henceforth, in our quantitative data analysis, we set the unmeasurable parameters 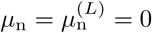 and subsume their uncertainties into an effective maximum generation *L*_e_. Finally, we invoke Eq. (4) to find the constraint

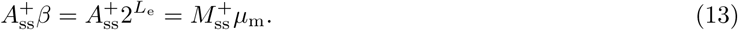

Since we can use the experimental value of 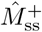 and *µ*_m_ has been reliably measured in the literature, Eq. (13) constrains 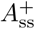 to *L*_e_.

After assigning values to parameters using Table 1 (setting *µ*_n_ = 0, *ω* = 0.16 and *µ*_m_ = 1), subsuming parameters into *L*_e_ (setting 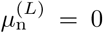), describing the configuration {*h_i_*} through *λ* and *C*_h_ (setting *µ*_h_ = 0), and applying the constraint 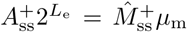, we are left with four effective model parameters *θ*_model_ = {*λ*, *C*_h_, *r*_n_, *L*_e_}. Here we have included *r*_n_ in the key model parameters since it is not reliably measured and the cell burst width is sensitive to *r*_n_. Once *L*_e_ is inferred, Eq. (13) can be used to find 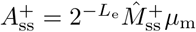.

### Statistical model

The total number of tags observed across all samples (obtained by summing up the observed numbers of *unique* tags over *J* samples) can be used as a lower bound on *C*_h_. Even though estimates for animal RQ5427 give *C*_h_ ~ 550 − 1100, the uncertainties *p*_h_, *K*_h_, and *H*(0) makes *λ* and *P*(*h*, *t*) difficult to quantify. Even if *P*(*h*, *t*) were known, it is unlikely that the drawn {*h_i_*} will accurately represent those in the monkey, especially when *λ* ≈ 1 and *P*(*h*) becomes extremely broad (the variance of *P*(*h*) approaches infinity). Thus we are motivated to find a statistical measure of the data that is insensitive to the exact configuration of {*h_i_*}. The goal is to study the statistical correlations between various features of *only* the outputs, which should be insensitive to the input configuration {*h_i_*} but still encode information about the differentiation dynamics.

Two such features commonly used to fit simulated *f_i_*(*t_j_*) to measured 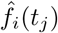 are the mean 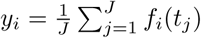 and the variance 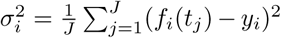. However, the small number of measurement time points *J* and the frequent disappearance of clones motivated us to propose an even more convenient statistic that is based on

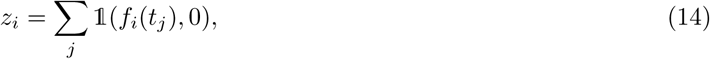

the number of absences across all samples of a clone rather than on *σ_i_*. Here, the indicator function 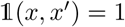 when *x* = *x’* and 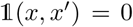 otherwise. In Appendix E in S1 Text, we illustrate alternatives such as data fitting based on *σ_i_* and on an autocorrelation function but also describe the statistical insights gained from using statistics of *z_i_*.

The level of correlation between 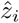 and 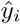 is measured by the average of 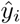 conditioned by their number of absences 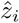 (dashed curve) in Fig. 4, where the distribution of the values of 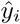 at each 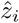 is clearly shown. To combine the correlated stochastic quantities *z_i_* and *y_i_* into a useful objective function, we take the expectation of *y_i_* over clones that have the same *z_i_*:

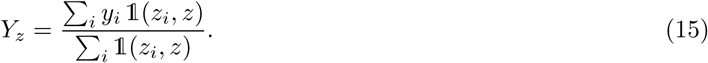

**Fig 4.**
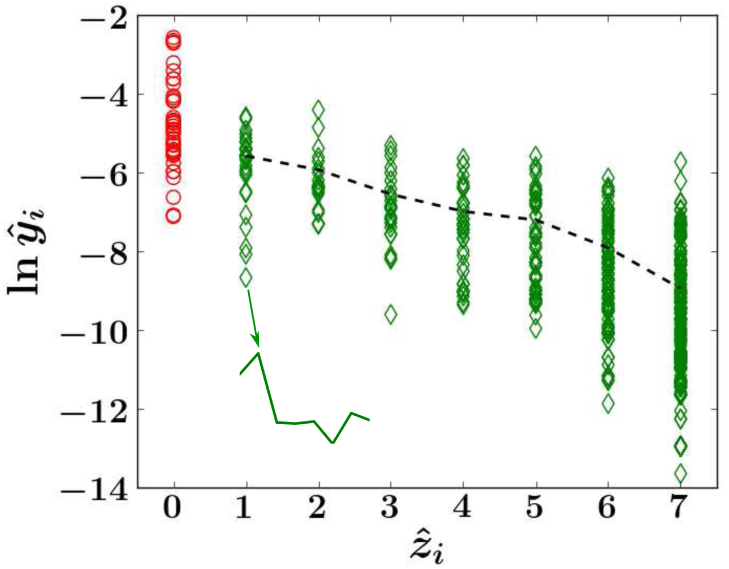
Scatterplot of clone trajectories of animal RQ5427 displayed in terms of ln 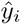, the log mean abundance of clone *i*, and 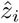, the number of samples in which clone *i* is undetected. The trajectory of each clone *i* is represented by a symbol located at a coordinate determined by its value of ln 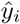 and 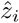. A trajectory of a clone that exhibits one absence within months 8 − 67 is shown in the inset. The first sample at month 2 is excluded because only long-term repopulating clones are considered. Clones that are absent in all eight samples are also excluded, so the largest number of absences considered for animal RQ5427 is 7. The dashed black line denotes 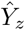, where 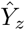 is the average of 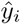 calculated over *i* within each bin of *z* as shown in Eq. (15). When later analyzing 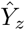, 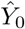 (red circles) is not included.

In case no simulated or data-derived trajectories *f_i_*(*t_j_*) exhibit exactly *z* absences, we set *Y_z_* = 0. We then determine *Y_z_* (*θ*_model_) from simulating our model and 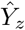 from experiment and use the mean squared error (MSE) between the two as the objective function:

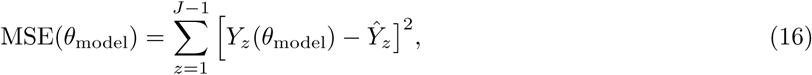

where *θ*_model_ = {*λ*, *C*_h_, *r*_n_, *L*_e_} and the sum is taken only over those *z* for which both data and simulations produce at least one clone (in practice, when searching for the best fitting *θ*_model_, we ensure at least 30 clones in each bin of *z*). Here *Y*_0_ is excluded from the MSE calculation because the *y_i_* values of clones that have *z_i_* = 0 are not constrained by the burstiness of the model and *Y*_0_ can be sensitive to the underlying configuration {*h_i_*} (see the Discussion and Appendix E in S1 Text).

We are now in a position to compare results of our model with experimental data. The general approach will be to choose a set of parameters, simulate the forward model (including sampling) to generate clone abundances {*f_i_*(*t_j_*)}, number of absences *z_i_*, and ultimately *Y_z_*(*θ*_model_), which is then compared to data-derived 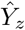. By minimizing Eq. (16) with respect to *θ*_model_, we obtain their least square estimates (LSE). A schematic of our workflow is shown in Fig. 5. We describe the details of the simulation of our model in Appendix F in S1 Text.

**Fig 5.**
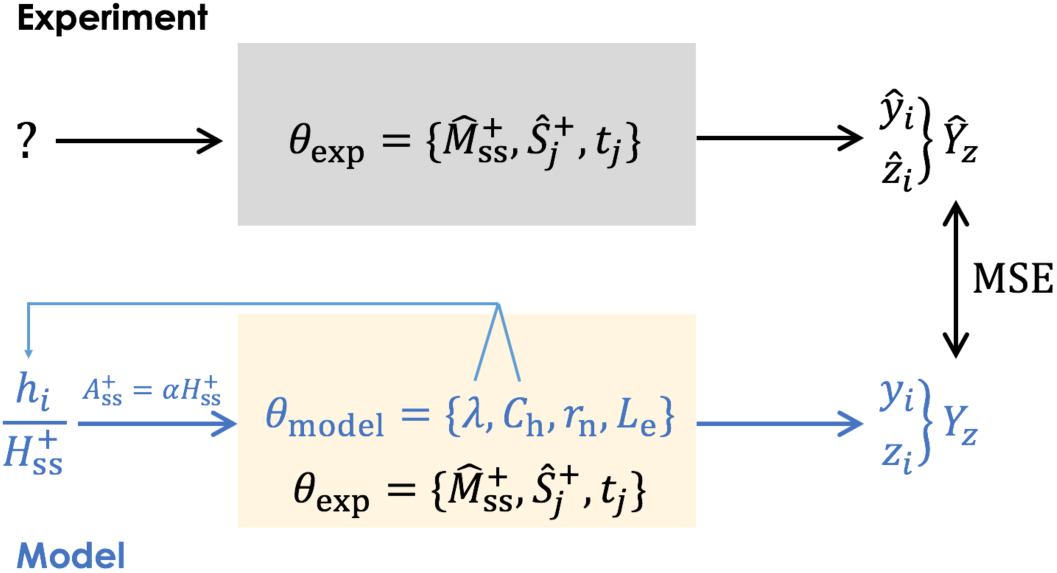
Workflow for comparing parameter-dependent simulated data with measured clone abundances. The initial input is the HSC clone distribution *P*(*h*), which is unknown and experimentally unmeasurable. Using known experimental parameters *θ*_exp_ and choosing model parameters *θ*_model_ the theoretical quantities *y_i_* and *z_i_* are computed by simulating the mechanistic model and the sampling. The corresponding 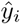 and 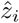 are extracted from data and the theoretical *Y_z_*(*θ*_model_) and the experimental 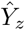 compared through the MSE defined in Eq. (16). The MSE is then minimized to find least squares estimates for *θ*_model_.

## Results

By implementing the protocol outlined in Fig. 5, we find a number of results including the shape of the MSE, least-squares-estimates (LSE) of the parameters, validation of the mechanistic model, and sensitivity analysis.

### Shape of the MSE function

For the range of values *r*_n_ ∊ [0.01, 10] and *L*_e_ ∊ [19, 28], the MSEs are fairly insensitive to *λ* ≥ 0.5 and 500 ≤ *C*_h_ ≤ 1000, but typically has lower values near *λ* ≈ 0.99 and *C*_h_ ≈ 500. Note that *C*_h_ ≈ 500 is close to the experimental estimate for animal RQ5427. Therefore, we fix *λ* = 0.99, *C*_h_ = 500 and minimize the MSE with respect to *r*_n_ and *L*_e_. For each {*r*_n_, *L*_e_} pair, simulation of the full model is repeated 200 times to generate 200 values of all (*y_i_*, *z_i_*) pairs, *Y_z_*(*λ* = 0.99, *C*_h_ = 500, *r*_n_, *L*_e_), and MSE(*λ* = 0.99, *C*_h_ = 500, *r*_n_, *L*_e_). The average values of these MSEs are plotted as a function of *r*_n_ and *L*_e_ in Fig. 6.

**Fig 6.**
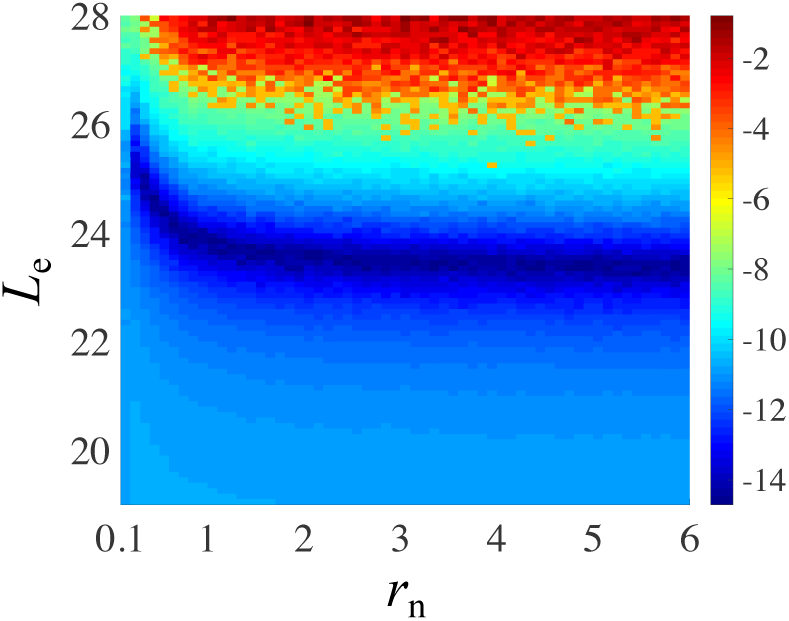
Dependence of the mean MSE defined in Eq. (16) on *r*_n_ and *L*_e_. For visualization purposes, we took the natural logarithms of MSE values and plotted them as a function of *L*_e_ and *r*_n_. Blue area denotes smaller MSE values, thus better fitting. This energy surface was generated by averaging over 200 simulations using *C*_h_= 500 and *λ* = 0.99.

We find that the minimum of the MSE is relatively insensitive to *L*_e_ for *r*_n_ ≳ 1. To interpret this result, note that *r*_n_ does not affect the absolute value of *β* according to Eq. (13), but it affects the typical time ~ *L*/*r*_n_+1/*ω* it takes for a generation-0 progenitor cell to form a mature granulocyte. When *r*_n_ < *µ*_m_, the proliferation of progenitors cannot “catch up” with the loss of granulocytes, resulting in a quickly vanishing burst in *m*_b_. A larger *L*_e_ would be required to compensate. When *r*_n_ ≫ *µ*_m_, the accumulation of *m_i_*(*t*) is much quicker than its loss so the burst size is relatively stable and 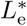 is not very sensitive to *r*_n_. Thus, the MSE objective function is fairly insensitive to *r*_n_ in its biologically meaningful value range.

### Least-squares estimates of *L*_e_ and 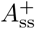 for animal RQ5427

To obtain an explicit best-fit value for *L*_e_ we fix *r*_n_ = 2.5 [12] (and *λ* = 0.99, *C*_h_ = 500) and varied *L*_e_ ∊ [19, 26]. The MSE objective function as *L*_e_ is varied is shown in Fig. 7(a). For *one* simulation at each chosen value of *L*_e_, we can construct the MSE and find an LSE for *L*_e_. Over 200 sets of simulations (for each chosen *L*_e_), we find the expected LSE value 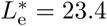, with a standard deviation of ±0.12. So in practice, the randomness across simulations is negligible. Upon applying the constraint in Eq. (13), the corresponding 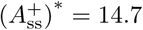. Substituting the LSE results into Eq. (11) yields a burst width of Δ*τ*_b_ ≈ 32 days, which is consistent to our assumption Δ*τ*_b_ ≪ Δ*t_j_* = 5 − 11 months. Fig. 7(b) shows how 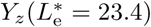 fits 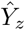.

**Fig 7.**
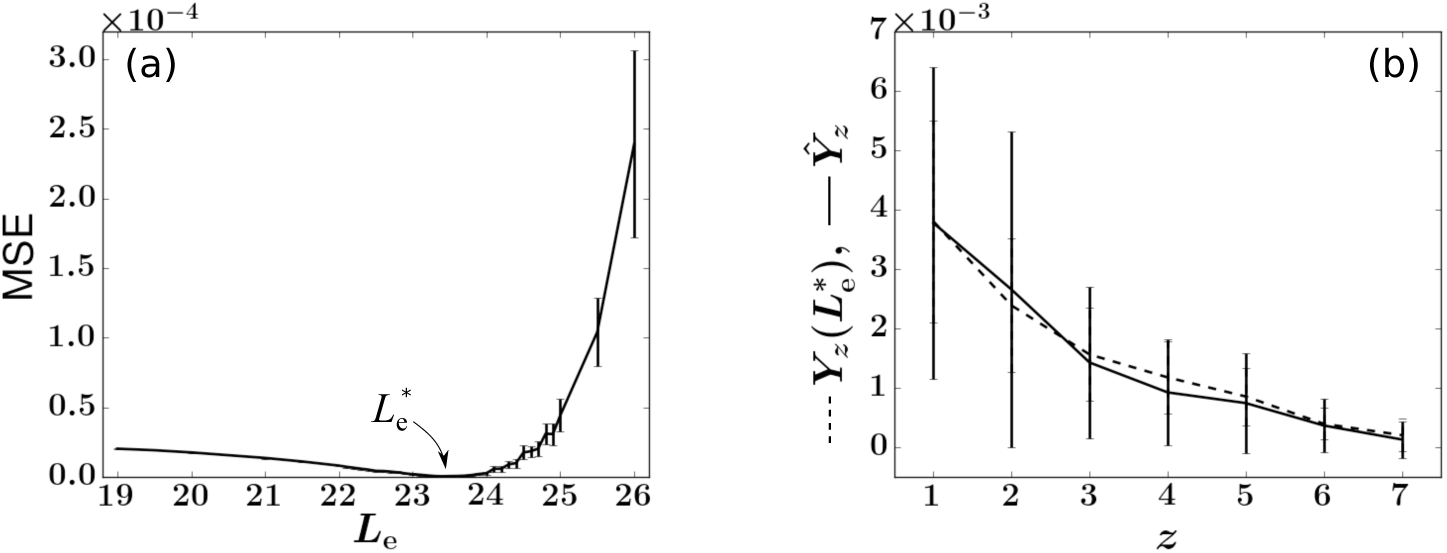
Finding the least squares estimate (LSE) 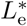 for animal RQ5427 by fitting the simulated *Y_z_* to the experimental 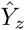. The values of (*λ*, *C*_h_, *r*_n_) are chosen to be (0.99, 500, 2.5). Simulations with {*h_i_*} set to 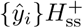 instead of drawing from *P*(*h*) generate similar results. (a) The LSE is 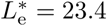. Averages and standard deviations (error bars) of the 200 MSEs are plotted. (b) Comparisons between the experimental (solid) 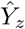 and simulated (dashed) *Y_z_* with fixed 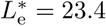. The error bars are determined by considering the standard deviation of the average abundances (*y_i_* or 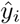) of all clones exhibiting *z* absences.

### Comparison of variability from simple sampling and best-fit model

We can check how our LSE result performs against the null hypothesis that clone size variations arise only from random sampling. An estimate of sampling-induced variability can be obtained by assuming a specific number of peripheral blood granulocytes of tag *i* and randomly drawing an experimentally determined fraction *ε*(*t_j_*) of peripheral blood cells. This is repeated *J* times from a constant peripheral pool {*m_i_*}. Each draw results in *si*(*t_j_*) cells of clone *i* in the simulated sample. Normalizing by *S^+^*(*t_j_*), the total number of tagged cells in the sample, we can define the rescaled mean abundance 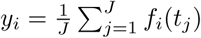 and the rescaled standard deviation 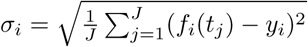 for each clone *i*.

The simulated quantities ln *y_i_* and *σ_i_* associated with each clone *i* and its experimental sampling fraction *ε*(*t_j_*) are indicated by the green triangles in Fig. 8(a). The corresponding values ln 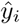 and 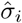 derived from the data shown in Fig. 1(b) are indicated by the blue dots. This simple heuristic test shows that the experimental fluctuations in clone abundances are significantly larger than that generated from random sampling alone and that additional mechanisms are responsible for the fluctuation of clone abundances in peripheral blood. Fig. 8(b) shows the fluctuations in clone abundances obtained from random sampling of fluctuating mature clones simulated from our model, using LSE parameter values. Here, the variability is a convolution of the fluctuations arising from intrinsic burstiness and from random sampling. The total variability fits those of the experimental data well except for several large-sized outlier clones.

**Fig 8.**
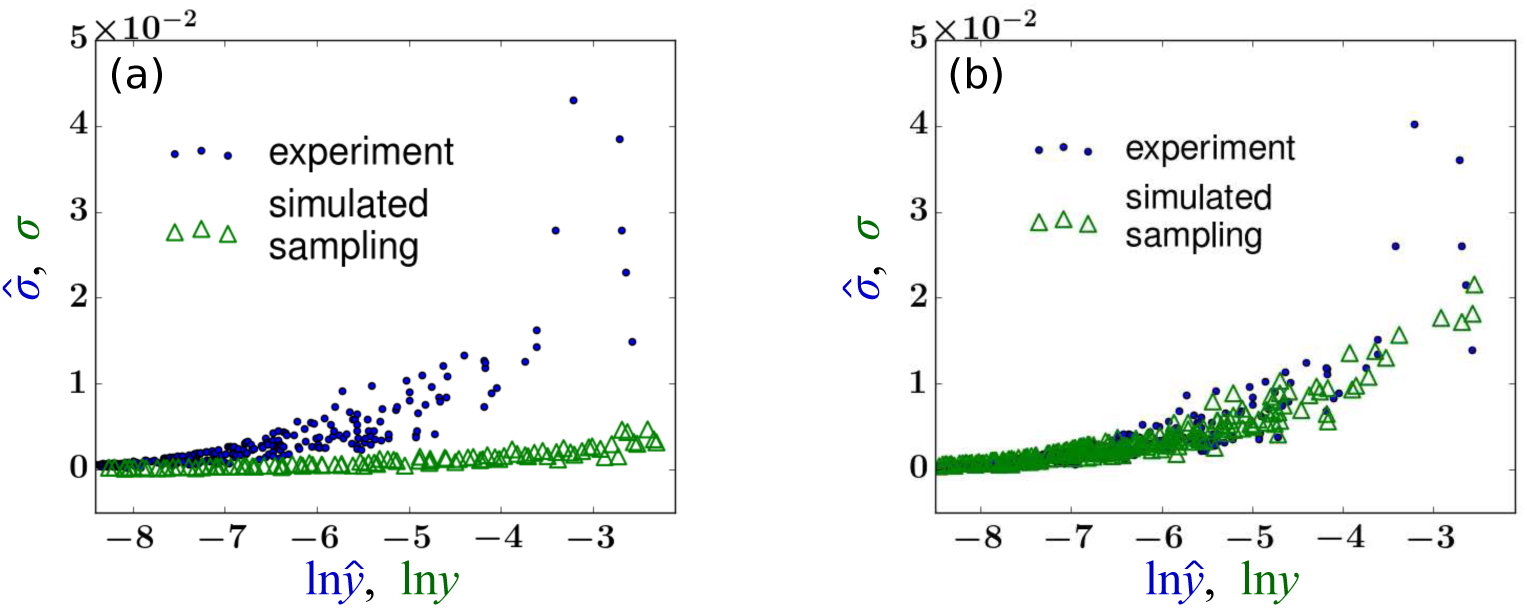
(a) A plot of the standard deviation 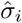 *vs*. the log of the mean 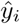, extracted from abundance data (blue dots). For comparison, clonal tags distributed within the peripheral blood cells were randomly sampled (with the same sampling fraction *ε*(*t_j_*) at times *t_j_* as in the experiment). The analogous quantity *σ_i_* shown by the green triangles indicate a much lower standard deviation for a given value of ln *y_i_*. This simple test implies that the clonal variability across time cannot be explained by random sampling. (b) The same test is performed after applying our model with the LSE parameter *L*_e_ = 23.4 (and the average of parameters listed in Table 1).

### Insensitivity of analysis to HSC configurations

We demonstrate the weak dependence of our least squares estimate to *λ*, the parameter controlling the shape of *P*(*h, t*), as shown in Fig. 9. For each *λ*, we sample a fixed number (*C*_h_ = 500) of *h_i_* from the theoretical distribution *P*(*h*, *t*) and let *L*_e_ vary between 19 and 28. We then simulate the model 200 times and find 200 MSEs at each value of *L*_e_ ∊ [19, 28]. The averages of the 200 MSE’s at each value of *L*_e_ are compared and the 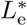 that corresponds to the minimal average MSE is selected. The selected 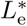 as a function of *λ* is plotted in Fig. 9(a). Fig. 9(b) shows the averages and standard deviations of MSE(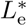) at each value of *λ*.

**Fig 9.**
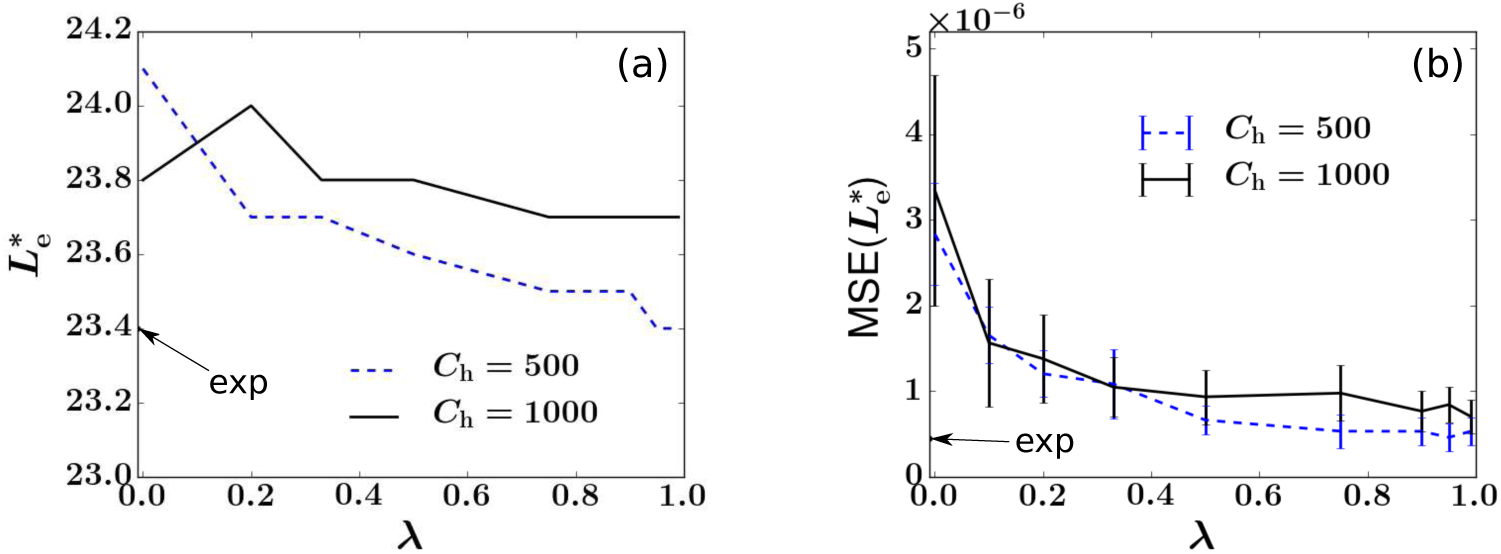
The LSE 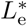 is insensitive to the geometric distribution factor *λ* > 0 and to *C*_h_ ≫ 1. This implies that for a wide range of values of *λ* and *C*_h_ the LSEs are insensitive to the HSC configuration {*h_i_*}. (a) 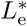 found at each value of *λ*. (b) Averages and standard deviations (error bars) of 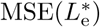 as a function of *λ.* The LSE and 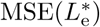 values associated with self-consistently using 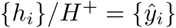 from experimental data are marked by arrows and “exp.”

We then repeat the simulations with *C*_h_ = 1000. These results together show that 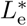 is insensitive to the distribution of *h_i_*. This insensitivity might be understood by noticing that *Y_z_* is defined as the *mean* of the values of *y_i_* that are associated with *z* absences (dashed curve in Fig. 4), and is not necessarily sensitive to how these values are distributed (vertically distributed markers at each value of *z* in Fig. 4). Instead, *Y_z_* encodes the intrinsic correlation between *y_i_* and *z_i_* and how much burstiness is transmitted to a clone’s *f_i_* (*t_j_*) from its *h_i_*.

To conclude, though it is generally impossible to recover the exact {*h_i_*} configuration, we find the HSC self-renewal-induced geometric distribution in Eq. (7) with factor *λ* ≥ 0.5 generates consistent comparisons with the sampled data.

### Data analysis and fitting for animals 2RC003 and RQ3570

The data from the three different monkeys vary in their numbers of tagged clones transplanted and the lengths of the experiments. For animal RQ5427/2RC003/RQ3570, there are 536/1371/442 clones that are detected at least once within 67/103/38 months. The fraction of cells in all tracked clones in animal RQ5427/2RC003/RQ3570 was approximated by the average fraction of cells that were EGFP+ marked over time, around 0.052/0.049/0.086 (the ratios between green square and blue triangle markers in Figs. 1(a), 10(a), and 11(a)), respectively. Figs. 10 and 11 also show the clone abundances, the MSE functions, and the statistics of *Y*(*z*).

**Fig 10.**
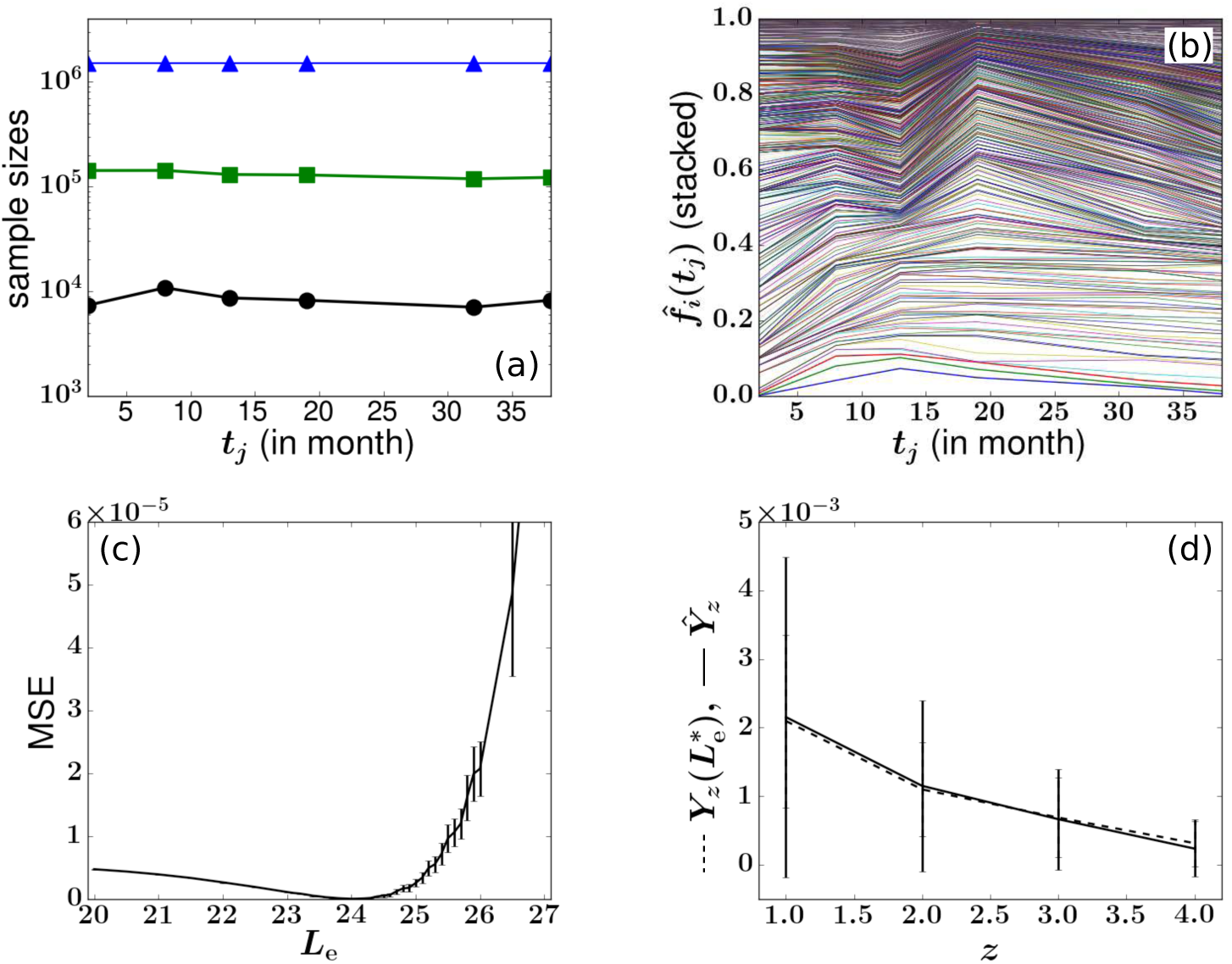
(a-b) Experimental data for animal 2RC003. (c) Difference between experimental 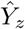 and simulated *Y_z_*(*L*_e_) as a function of *L*_e_. The values of hi’s are set to be equal to 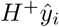 and the model was simulated 200 times at each value of *L*_e_. Other parameters are taken from Tables 1 and 2. The LSE 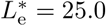 and 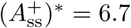. (d) Comparison of the optimal *Y_z_* to the experimental 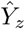.

**Table 2.**
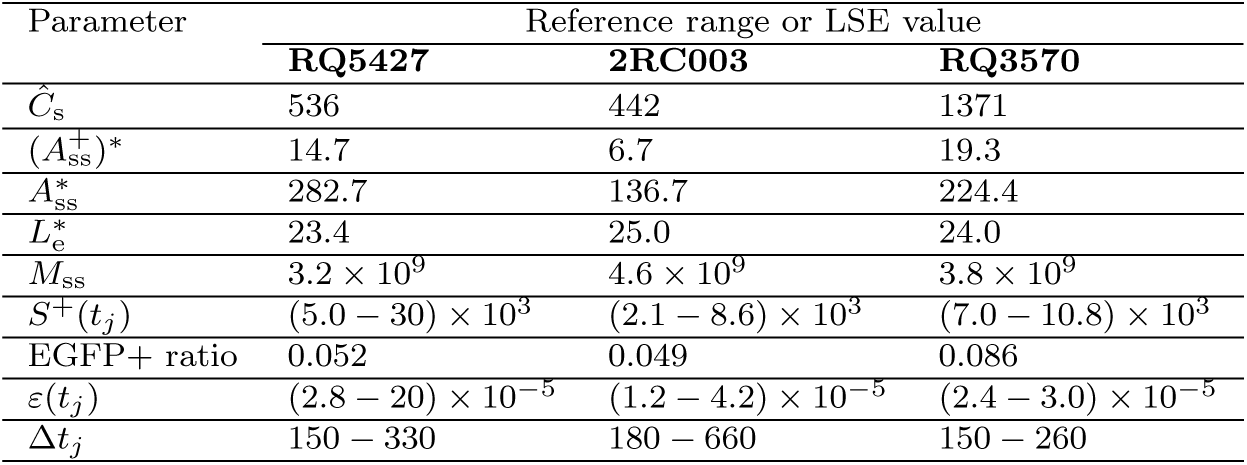
Summary of specific parameter values for monkeys 2RC003 and RQ3570 derived from experimental measurements [13] or obtained by calculations (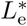 and 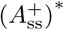).

| Parameter | Reference range or LSE value |  |  |
| --- | --- | --- | --- |
|  | RQ5427 | 2RC003 | RQ3570 |
| $\hat{C}_s$ | 536 | 442 | 1371 |
| $(A_{ss}^+)^*$ | 14.7 | 6.7 | 19.3 |
| $A_{ss}^*$ | 282.7 | 136.7 | 224.4 |
| $L_e^*$ | 23.4 | 25.0 | 24.0 |
| $M_{ss}$ | $3.2 \times 10^9$ | $4.6 \times 10^9$ | $3.8 \times 10^9$ |
| $S^+(t_j)$ | $(5.0 - 30) \times 10^3$ | $(2.1 - 8.6) \times 10^3$ | $(7.0 - 10.8) \times 10^3$ |
| EGFP+ ratio | 0.052 | 0.049 | 0.086 |
| $\varepsilon(t_j)$ | $(2.8 - 20) \times 10^{-5}$ | $(1.2 - 4.2) \times 10^{-5}$ | $(2.4 - 3.0) \times 10^{-5}$ |
| $\Delta t_j$ | 150 - 330 | 180 - 660 | 150 - 260 |

Despite differences among the animals and the large variability in the estimated values of *α* and *H*_ss_ individually reported in the literature [4,11,12], the estimates of 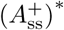 and 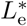 are rather similar across the three animals. For animal 2RC003, the optimal estimates are 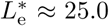, while for animal RQ3570, 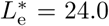. The corresponding estimates for *A*^∗^, after considering the constraint Eq. (13) and the EGFP+ ratios in Table 2, are 282.7, 136.7, and 224.4.

We also compared how the simulated LSE 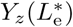 fits the experimental 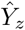 for all three animals. Note that for each specific *z*, the value of *Y_z_* is the conditional mean of the values of *y_i_* for which clones *i* exhibits exactly *z* absences. To evaluate the “quality” of fitting 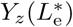 to 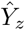, one can directly perform a two-sample t-test between the two sets of values 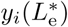 and 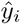 that contribute to each value of *z*. The group of 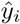 values corresponding to each value of *z* is shown by the vertical cluster of diamonds in Fig. 4, while the corresponding set of values of 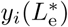 are generated by simulations. For each *z* value, we performed 200 simulations and collected the values 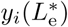 of all clones *i* that exhibit *z* absences and contribute to 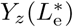. We ensured at least 30 values of 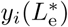 for each *z* and performed the t-test with the measured set 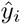 containing clones that exhibit the same number of absences *z*. Performing this t-test for all 1 ≤ *z* ≤ *J* – 2, we generate 200(*J* – 2) p-values. Data for *z* = *J* – 1 is too noisy and was not included. A p-value *p <* 0.05 would indicate that the two sample means *Y_z_*(*L*^∗^) and 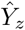 are not likely to be considered identical; if instead *p* ≥ 0.05, the null hypothesis of equal sample means cannot be rejected. For animals RQ5427 and RQ3570, 93.5% and 99.4% of the p-values are larger than 0.05, while for animal 2RC003, the fraction is 76.6%. This result is consistent with the eroded fitting quality with increasing experimental time, where the slow decrease in the number *C*_h_(*t*) of HSC clones in animal 2RC003 cannot be neglected. As evident from Fig. 10(a), several clones start to dominate after month 64; this coarsening phenomenon is not evident in the data of the other two monkeys. Animal RQ3570 was sacrificed at month 38 so no obvious coarsening is observed and no clones strongly dominate (see Fig. 11). A summary of the parameters and fitting results for all animals is given in Table 2.

**Fig 11.**
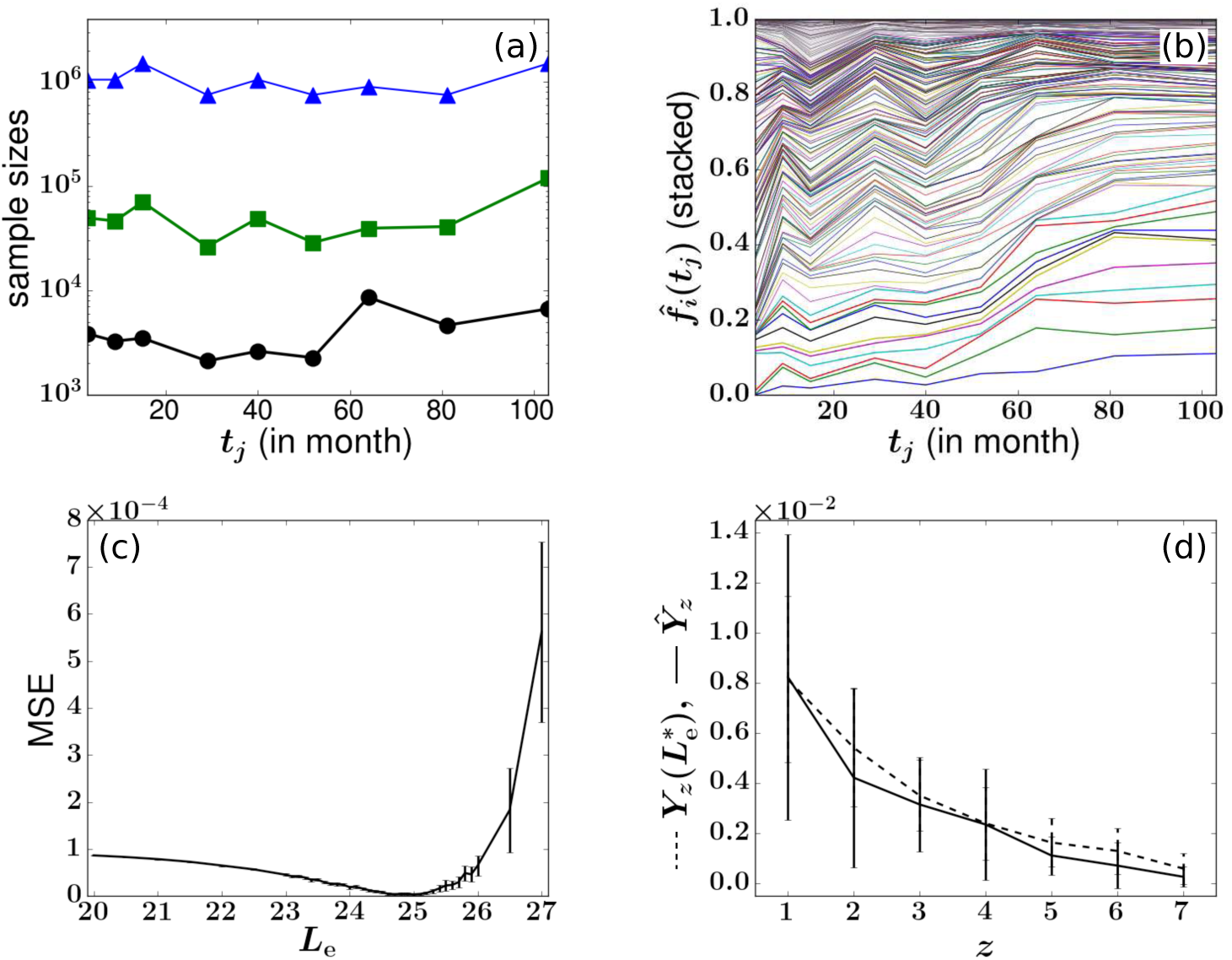
Experimental data (a-b) and fitting results (c-d) for animal RQ3570. The values of *h_i_*’s are set to be equal to 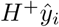. Other parameters are taken from Tables 1 and 2. The LSE fitting results are 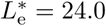 and 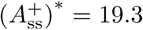.

## Discussion

In this study, we analyzed a decade-long clonal tracking experiment in rhesus macaques and developed mechanistic and statistical models that helped us understand two salient features of clone abundance data: the heterogeneous (nonuniform) distribution of clone sizes and the temporal fluctuation of clone sizes. Below, we further discuss the implications of our results, the structure of our mechanistic model, and the potential effects of including additional biological processes.

### Comparison to previous studies

The long-term clonal tracking data we analyzed were generated from a huge number of initially tagged HSPCs (*C*_h_(0) ~ 10^6^ − 10^7^) [13], a large number of observed clones (*C*_s_ ~ 10^2^ − 10^3^), small numbers of sequenced cells (10^3^ − 10^4^), and infrequent sampling. This presents significant challenges to the modeling and analysis over previous studies that mostly focused on one or a few clones [5,15,17,18].

In a previous analysis, Goyal et al. [4] aggregated the clone abundance data across *all* mature cell types and studied the distribution of the *number* of clones of specific size. At each time point, they ordered the clones according to their sizes. Thus, the ordering can change across samples as some clones expand while others diminish. They found that the cumulative clone number distribution (defined as the number of clones of a specific size or less) of the size-ordered clones become stationary as soon as a few months after transplantation. They proposed a neutral birth-death description of progenitor cells and fitted the *expected* value of clone counts in each sample by assuming 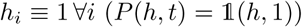 and tuning parameters in the downstream progenitor and mature cell compartments. By focusing on aggregate clone counts, this study could not distinguish the dynamics of individual clones nor could it predict the persistence of clone sizes over time. Since individual clone sizes (*h_i_*, *n_i_*, *m_i_*, *s_i_* of the same tag *i*) were not tracked, mechanisms driving the dynamics, and in particular, the variability and fluctuations of *individual* clone sizes that drive disappearances and reappearances, remain unresolved [4].

In our model, heterogeneity of clone sizes is explicitly generated by stochastic HSC self-renewal of cells of each tag and extinctions and resurrections arise from a generation-limited progenitor proliferation assumption. We infer model parameters as listed in Table 2. Combining the results with previous experimental and theoretical estimates of *H*_ss_ ≈ 1.1 × 10^4^−2.2 × 10^4^ [4,48] results in *α* = 0.0045−0.027, slightly larger than, but still consistent with, the estimates *α* = 0.0013 − 0.009 by Shepherd *et al.* [11]. Previous studies that modeled total peripheral blood population estimated *α* ≈ 0.022 and *H*_ss_ ≈ 1.1 × 10^6^/kg for dog and *α* ≈ 0.044 and *H*_ss_ ≈ 1.1 × 10^6^/kg for human [12]. These estimates yield a value of *αH*_ss_ about 10^2^ − 10^3^ times greater than ours, which is nonetheless consistent with our steady-state constraint Eq. (13) because they assumed a much smaller *L* ≈ 15 − 18 for dog and 16 − 21 for human. This difference in the estimates of *L* may be partially attributed to the transplant conditions under which the rhesus macaque experiments were performed [13]. Alternative model assumptions and differing values of other parameters may also contribute to this difference. For example, the extremely large value of *H*_ss_ ≈ 10^7^ used in [34] will naturally decrease their estimate for 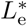 relative to that of our analysis.

### Model structure, sensitivity to parameters, and cellular heterogeneity

Uncertainties in values of parameters such as *µ*_h_, *p*_h_, *K*_h_, and other factors that tune the symmetric-asymmetric modes of HSC differentiation or involve HSC activation processes [49] will impart uncertainty in determining *P*(*h*) and {*h_i_*}. We have assumed *P*(*h*) satisfies a master equation and depends only two effective parameters *λ* and *C*_h_. However, we have demonstrated that the statistical properties of *Y_z_* are quite insensitive to the upstream configuration {*h_i_*} and hence to *λ* and *C*_h_ for a wide range of their values (see Fig. 9). In other words, very little information in {*h_i_*} is retained in the sampled abundances 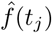 after HSCs differentiate and trigger random bursty peripheral blood cell population dynamics.

Another feature we have ignored in our neutral model is cellular heterogeneity such as tag-dependent differentiation, proliferation, and death rates. Cellular heterogeneity in HSC differentiation rates could be described by different *α_i_* for each clone *i* and the total differentiation rate would be 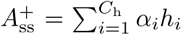. Differences in *α_i_* can be subsumed into a modified configuration {*h_i_*} which, as we have seen, does not strongly influence our parameter estimation based on the *Y_z_* statistics. Thus, given the available data and how information is lost along the stages of hematopoiesis and sampling, the present quasi-steady state analyses cannot resolve heterogeneity across HSC clones.

We have not investigated how cellular heterogeneity in progenitor and mature cells would affect our results, but clone-dependences in their birth and death rates could affect sizes and durations of population bursts and quantitatively affect our analysis. However, unless the statistics of inter-burst times are highly variable across clones, we do not expect cellular heterogeneity to qualitatively affect our conclusions.

Changing downstream parameters such as *µ*_m_ or invoking alternative mechanisms of terminal differentiation (see Appendix B in S1 Text) can affect the shape of clonal bursts. We show in Appendix D in S1 Text that these effects can be subsumed into the effective maximum progenitor generation *L*_e_. We have performed additional simulations to confirm that changing *µ*_m_ = 2 will not influence the fitting of 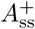 but but increases 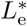 by one. In other words, inference of 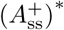 is robust against many upstream and downstream parameters, indicating that the intrinsic clone size fluctuations observed in the experimental data strongly constrain the total rate of HSC differentiation. On the other hand, uncovering the actual maximal generation *L\** from 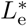 is possible only when uncertainties in these other parameters are resolved.

### Clonal stability *vs* clonal succession

Our model reduction was based on the separation of timescales of the slow HSC dynamics and the fast clonal aging dynamics. Since HSC clone sizes vary extremely slowly for primates (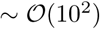 months), we ignored the homeostatic births/deaths of HSCs when fitting the temporal clonal variations. This is partially justified by visual inspection of Figs 1(b), 10(b), and 11(b) that no significant variations of large clones’ abundances is observed before 60 months. Instead, the random intermittent HSC differentiation events induce relatively short (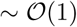 months) bursts of granulopoietic progeny that contribute strongly to temporal fluctuations of clone sizes. Such behavior is consistent to the “clonal stability” hypothesis [50–52], which assumes that a fixed group of HSCs randomly contribute to an organism’s blood production at all times.

The alternative hypothesis of “clonal succession” [16, 53, 54] assumes that different groups of HSCs are sequentially recruited to the blood production at different times. This hypothesis would only be consistent with our model under a different set of parameters where HSCs self-renew/die at a rate comparable to that of Δ*τ*_b_, the duration of a granulocyte burst. For example, murine HSC turnover rates *µ*_h_ are hypothesized to be 10-fold higher than those in primates while the clonal aging dynamics (and its timescale Δ*τ*_b_) are relatively conserved across species [55]. According to our result in Appendix B in S1 Text, such a 10-fold increase in HSC death rate would lead to a 10-fold increase in HSC clone extinction rate, bringing the lifespans of HSC clones closer to the (progenitor) clonal aging timescale Δ*τ*_b_. This interpretation is consistent with the fact that hematopoiesis in large primates have been described in terms of “clonal stability” while hematopoiesis in mice have been described in terms of “clonal succession” [16,50–54]. We thus predict that with even longer tracking (> 100 months), the “clonal succession” mechanism could also be significant in primates.

### Summary and future directions

In summary, we have built mechanistic and statistical models that enable the quantitative analysis of noisy and infrequent clonal tracking data. We focused on the huge temporal variability observed in the sampled clone abundances, defined a robust statistical measure *Y_z_* of sample-to-sample clone size variability through the number of clonal disappearances. Of course there is a nearly endless list of details such cellular heterogeneity and more complex biology that we did not include, but given the noisy data, we propose and quantify the simplest explanation for the observed heterogeneous clone abundances and the temporal “extinctions and resurrections.” The key ingredients in our mechanistic model are HSC self-renewal (quantified by the effective parameter *λ*), intermittent HSC differentiation (quantified by the parameter 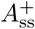), and an effective maximum progenitor generation (quantified by the effective parameter *L*_e_). Although we cannot fully resolve *λ* from data, the obvious mismatch between experiment and our model when *λ* is small shows that a certain level of HSC clone-size heterogeneity (larger *λ* ≈ 1) is necessary to match the sampled data. Similarly, we cannot fully resolve *α* and 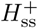, but their product, the total tagged HSC differentiation rate 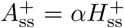, is one of the key parameters constrained by our modeling. By minimizing an objective function of *Y_z_* over effective model parameters, we found LSE values 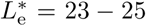 and 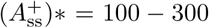 for the three rhesus macaques. These quantities could not be inferred from the total, more static cell populations. These results also imply that true dynamical changes in 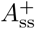 and *L*_e_ could be masked by the intrinsically bursty dynamics of each clone, but provide a framework for future study into extrinsic perturbations.

Our analysis provides insight into which variables and experimental conditions parameter inference is most sensitive to, possibly guiding the design of future experiments. The approach and models can also be readily extended to quantify white blood cells of other types. For example, the mechanistic model can be directly applied to monocytes since they also have relatively simple dynamics and do not proliferate in the periphery [56]. Peripheral lymphocytes, however, would require additional experimental information because their populations are more sensitive to the state of the animal and can homeostatically proliferate [38].

## Acknowledgements

This work was supported by grants from the NIH (R01AI110297 and K99HL116234), the California Institute of Regenerative Medicine (TRX-01431), the UCLA AIDS Institute/ Center for AIDS Research (AI28697), the Army Research Office (W911NF-14-1-0472), and the NSF (DMS-1516675). SX acknowledges support from the Chinese Scholarship Council (no. 201408020004). The authors also wish to thank anonymous reviewers whose detailed suggestions helped improve the manuscript.

